# The conflict negativity: A neural correlate of value conflict and indecision during financial decision making

**DOI:** 10.1101/174136

**Authors:** Gesa-K. Petersen, Blair Saunders, Michael Inzlicht

## Abstract

Individuals struggle when making financial decisions, sometimes preferring to avoid difficult decisions and receive lower future rewards over actively deliberating between options of similar value. Here, we examine how conflict deriving from objective and subjective value characteristics of stocks, as well as the behavioural and phenomenological correlates of decision conflict, are accompanied by variation in a thus far understudied ERP component, the conflict negativity (CN). In a novel EEG paradigm (N = 53), we simulated a financial decision situation in which participants made incentivized choices between different, sometimes conflicting, stock options. Our results indicate that participants become slower, more undecided, and less pleased, when choosing between similar options compared to choices in which one option clearly outweighs the other. This effect even held when participants chose between two objectively good alternatives. We further provide preliminary evidence that the CN, a negative-going ERP recorded over the medial prefrontal cortex, is not only sensitive to decision conflict, but also predicts behavioural indecision. What is more, subjective value characteristics of stocks, impressions based on brand perception of the stock options, modestly influenced affective and behavioural reactions over and above objective stock characteristics. While our results are at odds with assumptions made by classic economic theory, they may serve as one out of several indicators as to why private investors seem to avoid financial decisions.

---

Decisions are not easy, and this is especially true when it comes to our finances. Economic research suggests that people avoid making even relatively simple choices (cf. Johnson & Goldstein, 2003; Madrian & Shea, 2011; Thaler & Benartzi, 2004). Perhaps as a result of this, levels of trading activity are low, with one in five individuals with brokerage accounts making no trades at all (“Shareownership 2000”, 2002), a phenomenon commonly known as investor inertia. Planning and saving for retirement, for example, is a form of investment decision that affects almost all individuals. Yet, individuals often avoid difficult decisions in this domain and, in turn, make suboptimal provisions for their future financial security (Madrian & Shea, 2011). But what makes financial decisions, with seemingly rewarding outcomes, so aversive and avoidable? Here we explore this question, examining the neural, behavioural, and phenomenological correlates of indecision during value-guided choices in a simulated stock-market.

## Difficulty and discomfort during decision making

Preference-based decisions in scenarios with uncertain outcomes are often beset by indecision, anticipated regret, and unpleasant experiences—even when people choose between two objectively good alternatives (cf. Anderson, 2003; Richard, Van der Pilgt & De Vries, 1996; Schwarz, 2000; Zeelenberg, 1999). In one sense, it seems counterintuitive that these decisions would give rise to aversive experiences. Positive emotion might be expected in situations where people choose between two equally promising options. Philosophical perspectives of free-will, however, have proposed that paradoxes are prevalent during seemingly rational decision making. Aristotle, for example, stated that a person, “…exceedingly hungry and thirsty, and both equally, yet being equidistant from food and drink, is therefore bound to stay where he is…” (350 BC, trans. 1922, book II, part 13). As captured in the quotation, a purely rational decision making process is stymied when the individual is forced to deliberate between two equally appealing options.

While the allegory of the hungry and thirsty man stalling when equally situated between food and drink might sound overstated, modern accounts of affect and motivation suggest that indecision and choice conflict do provoke unpleasant experiences (Festinger, 1962; Proulx, Inzlicht, & Harmon-Jones, 2012; Shenhav & Buckner, 2014). Generally speaking, choice conflict in this context is defined as a state in which incompatible response tendencies are simultaneously active (e.g. Botvinick, Braver, Barch, Carter, & Cohen, 2001; Nakao et al., 2010), such as when two options in a choice dilemma are simultaneously equally appealing to an individual. With regards to occupational choices, neuroscience research using ERPs has demonstrated that decisions involving high conflict (i.e., when two or more response options are similarly valued) leads to higher levels of neurophysiological reactivity and slower response times than low conflict decisions in the same domain (Nakao et al., 2010; Nakao, Bai, Nashiwa, & Northoff, 2013).

## The conflict negativity

In the current study we explored the integration of the neurophysiological and affective correlates of effortful financial decision making using an ERP termed the conflict negativity (CN; Nakao et al., 2010; 2013). The CN (see figure 1) is a negative going ERP that often starts before the response, and typically peaks 50-100 ms after a response at frontal/central midline electrode sites (Di Domenico, Le, Liu, Ayaz, & Fournier, 2016; Lin, Saunders, Hutcherson, & Inzlicht, 2017; Nakao et al., 2010, 2013), and has been source localised to dipole generators within the anterior cingulate cortex (ACC; Di Domenico et al., 2016). EEG time-frequency analyses have further indicated that the conflict negativity largely reflects increased theta-band power at electrode sites over the medial prefrontal cortex (e.g., Cavanagh, Figueroa, Cohen, & Frank, 2011; Lin et al., 2017).

**Figure 1.**
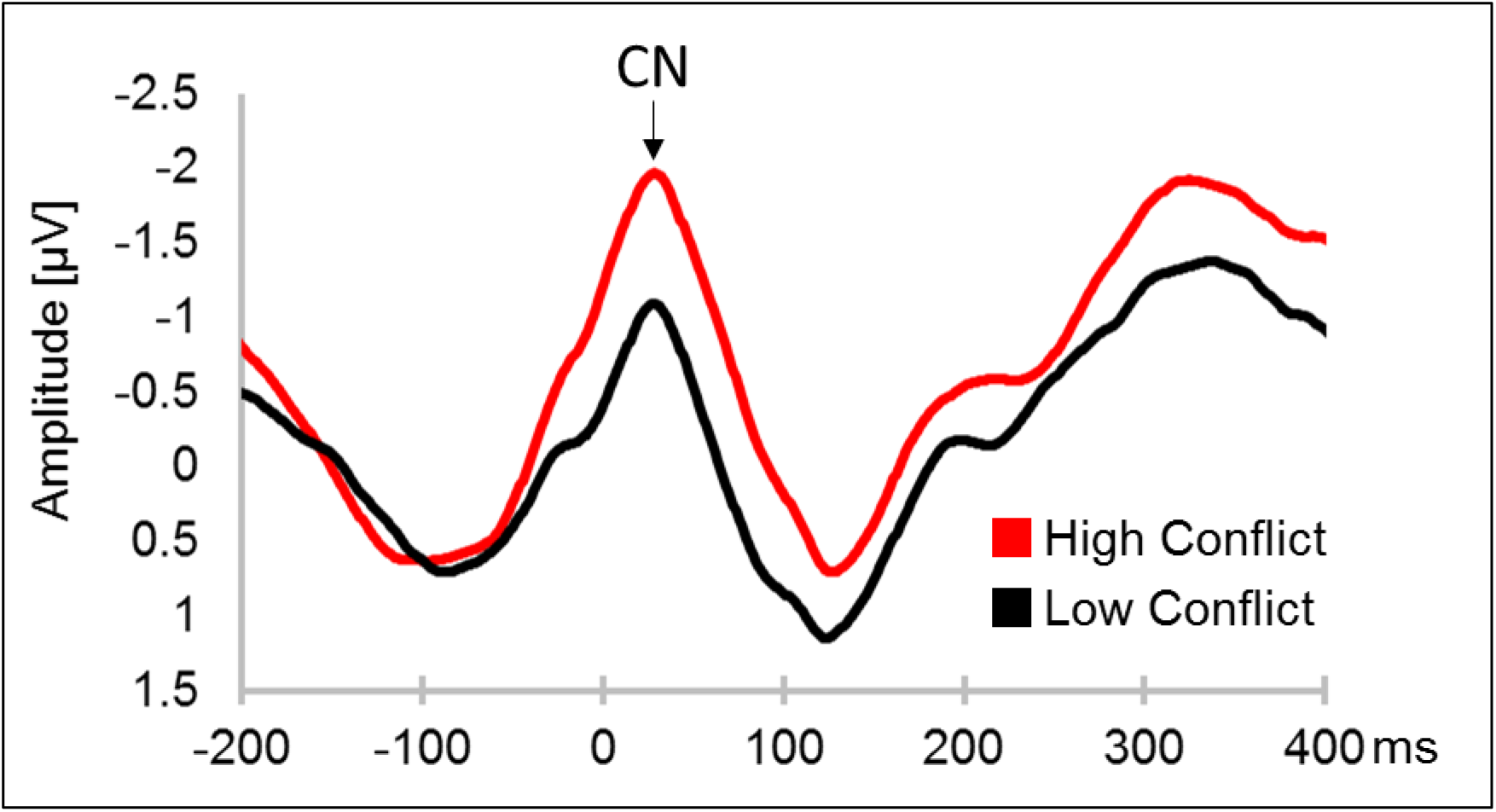
ERP waveforms depicting the conflict negativity during for high conflict and low-conflict decisions in a temporal discounting paradigm (adapted from Lin et al., 2017).

The psychophysiological profile of the CN suggests that the component very likely belongs to a family of performance monitoring related midline ERPs, including the error-related negativity (ERN), correct-related negativity (CRN), feedback negativity (FN), and N2 (cf., Holroyd & Coles, 2002; Yeung, Botvinick, & Cohen, 2005). While further investigation and source localisation are clearly warranted, it is reasonable to suspect that the CN, ERN, N2, and CRN reflect the activity of a common frontocentral monitoring system (see Cavanagh, Zambrano-Vazquez, & Allen, 2011). These ERPs have also been functionally related to similar psychological processes. More specifically, these frontocentral ERPs are thought to reflect a process that monitors for challenges to successful performance, such as conflict, and signals to other brain areas the need for adaptive adjustments in ongoing performance (Cavanagh & Frank, 2014; Kerns et al., 2004). One ongoing controversy relating to these ERPs is whether they are specific to conflict (e.g., Yeung et al., 2004), or whether they are sensitive to a wider range of control-relevant events such as error-likelihood (Brown & Braver, 2005), reward-prediction error (Holroyd & Coles, 2002), or surprise (Lin et al., 2017).

The CN perhaps bears closest resemblance to the CRN (Vidal, Hasbroucq, Grapperon, & Bonnet, 2000) that presents as a small negative deflection 0-100 ms after accurate performance, and is sensitive to variation in conflicts arising during executive functioning tasks such as the flanker paradigm (Bartholow et al., 2005). In addition to the objective conflict that arises during conflict control tasks, however, the CN also appears to be sensitive to conflicts in subjective, value based decision making. That is, the CRN emerges on tasks when participants make objectively correct responses (e.g., central arrow pointing left with participant indicating left), whereas the CN appears even when there is no objectively correct response (e.g., when participant prefers being a professor over a dentist). Supporting this idea, recent studies have indicated that the CN tracks subjective conflicts that arise when considering personal preferences for delayed rewards (Lin et al., 2017) or personal preferences during occupational choice (Di Dominico et al., 2016; Nakao et al., 2010). This functional characteristic of the CN means that it might be used to investigate conflict monitoring across a range of decision making domains that bear closer resemblance to the types of uncertain, subjective decisions that are common in everyday life. The sensitivity to subjective choice distinguishes the CN within this range of performance monitoring ERPs and motivates us to retain the distinct nomenclature. Furthermore, the name “*correct* related negativity” is something of a misnomer in relation to value-based decision making scenarios in which there is no objectively correct answer.

Relative to other performance monitoring ERPs (i.e., ERN, FRN, and N2) the CN (and CRN) is relatively understudied. Consequently, a key goal of the current work was to further explore the functional significance of the CN by testing the ERP’s associations with the behavioural and phenomenological correlates of financial decision making.

## Financial decision making

Financial decision making provides a useful domain in which to test the functional significance of the CN because the monetary value of choice options equalizes and objectivises the value parameters. Interestingly, with regards to normative economic theories (see Schoemaker, 1982; Fehr & Hoff, 2011), facing objectively equal options should induce no conflict or indecision at all. Furthermore, the slightest indication of one option outperforming the other—even by a single cent—should guide the rational decision maker without conflict. As a logical conclusion, financial decisions should be easy in both cases when available options are of equal value and when one option is clearly better than the other.

Somehow contradicting the implied ease of economic decisions, it was shown that a substantial fraction of employees in a large U.S. corporation avoided decisions on their retirement savings and simply accepted the suggested default setting (Madrian & Shea, 2011). The shown inertia may be a result of trying to avoid negative arousal. In one fMRI study, conflict between highly valuable consumer goods (e.g., selecting between a digital camera and an ipod) was associated with increased anxiety relative to choices involving one less desirable product options (Shenhav & Buckner, 2014). Furthermore, this anxiety co-varied with canonical conflict-related neural activity in the anterior cingulate cortex—a brain region associated with both negative affect and conflict monitoring (Shackman et al., 2011). In contrast to classic economic theories, these findings suggest that decisions between high value options are both conflicting and aversive—even when a desirable outcome is guaranteed.

Classic economic theories also imply that if options are equal, and choosing one or the other leads to an equivalent outcome, subjective valuations such as liking should not matter at all.

However, findings in behavioural finance (e.g., Madrian & Shea, 2011), in addition to the neuroscience of subjective decision making (e.g., Shenhav & Buckner, 2014), suggest that decisions are not made in this straightforward and rational manner. Real world investors consider subjective information next to economic factors when making decisions. For example, social information about mutual fund managers such as their ethnicity (Kumar, Niessen-Ruenzi, & Spalt, 2015) or gender (Niessen-Ruenzi & Ruenzi, 2015) influences fund inflows when controlling for objective characteristics such as risk and return. Thus, subjective preferences seem to influence financial decisions over and above economic factors, perhaps partially explaining why seemingly straightforward financial decisions can become beset with conflict and indecision.

In sum, though normative economic models suggest that no conflict should arise when making investment decisions, we propose, based on recent empirical economic data and the affective neuroscience of decision making, that investors will experience conflict and unpleasant emotion when making investment decisions between objectively equal options. We further suggest that subjective preferences matter over and above objective characteristics. Critically, we examined within-person affective, behavioural, and neurophysiological responses to financial decision conflict.

## Current study

Based on our propositions, we tested four general hypotheses regarding the role of conflict in financial decision making. First, we expect that decisions between equally valuable options will generate an aversive conflict state characterised by higher levels of behavioural indecision, increased amplitude of the CN (c.f., Nakao et al., 2010; Nakao et al., 2013), reduced positive affect, and increased anxiety relative to options with one clearly preferable option. Second, we tested if these conflict effects are amplified by expected value that is if conflict effects are larger when higher rewards are at stake. Third, we predicted that within-participant variation in the CN—as a neural correlate of conflict monitoring—would correlate with within-person variation in behavioural indecision, positive affect, and anxiety. In short, the magnitude of neural reactivity to conflict will go hand-in-hand with the behavioural and phenomenological correlates of decision difficulty. Fourth, we tested if subjective preferences for available investment options correlate with the neural, behavioural, and phenomenological correlates of conflict over and above the objective characteristics of the stock options.

To provide an ecologically valid environment while thoroughly testing these hypotheses in a controlled setting, we designed a stock-broker game. Like in a real stock-market, participants could earn money based on the risk and return characteristics of the investment options. Moreover, we used existing company names so that we had a genuine source for participants’ subjective preferences, and, in addition, so that decisions appeared more realistic.

## Method

### Transparent Reporting

Sample size was determined before data-collection commenced. The study was not pre-registered. All data, analysis scripts, and study materials are available on our project page on the Open Science Framework (https://osf.io/7tqj3/). All exclusions are reported in the following results method and results sections, with the sample sizes reflecting the net number of participants included after exclusion.

### Participants and design

The study consisted of two parts: An online survey to most notably elicit participants’ subjective preferences on the companies used in the stock broker game, and a laboratory experiment to actually play the stock broker game and meanwhile measure participants’ behavioural, subjective and neurophysiological conflict reactions. Altogether 56 undergraduate participants from the University of Toronto Scarborough took part in return for course credit. The advertisement to the participant pool offered the chance to play a stock-broker game, for which participants were compensated with course credit; they were also given the opportunity to earn a bonus cash prize. A power analysis using G*Power 3 (Faul, Erdfelder, Lang, & Buchner, 2007) testing a within–between interaction with small effect sizes (f=0.15) in a mixed model suggested collecting 50 (80 % power) to 64 (90% power) participants. We thus aimed to collect close to 60 participants and did not conduct hypothesis testing before termination of data collection. Three participants were excluded from the sample because they did not understand the experimental task (for further details see “procedure – learning phase”) leaving 53 participants. EEG analyses were conducted with a minimum of four trials per bin (please find results for three and five trials per bin in the supplemental online material: https://osf.io/mcf34/). 5 participants from the laboratory could not be matched with data from the online survey, so that data from these participants could not be used for combined analyses. Overall, participants were on average 19.17 years old (SD = 2.01 years, 90% CI [17, 22]), lived for 12.22 years in Canada (SD = 7.25 years, 90% CI [1, 21]) and 52.08% of all participants were female.

Each experimental session lasted ∼90 minutes in the laboratory plus 15 minutes for the online pre-test questionnaire. The experiment was approved by the research ethics board at the University of Toronto.

### Procedure

#### Online survey

Participants first completed an online survey at least 24 hours (M = 6.02 days, SD = 5.80 days, 90% CI [4.67, 7.37]) prior to the start of their laboratory session. The goal of this online survey was to elicit each participant’s subjective impressions of eight well-known companies from four sectors (technology, finance, retail, and motor manufacturing). These companies were used later for the virtual stock-broker game in the laboratory. In the online survey, participants first bid a price they would be willing to pay for a single share in each of the eight companies, based on the Becker-DeGroot-Marschak method (BDM; Becker, deGroot, & Marschak, 1964). Deviating slightly from the classic BDM design, participants were given a CAD 100 budget (instead of not providing any budget at all) and had to decide how much to invest by freely allocating a dollar amount that they would pay to receive a stock in each company. We changed this part in the design as we considered it to be a better way to approximate relative subjective values. Participants were told that their bid would have real consequences in the laboratory: “…indicate how much you would be willing to pay for these shares from your budget without knowing the actual stock-market value. If your bid for a share is equal or higher than the share’s real stock-market price, you will have purchased this share, if it is lower than the price you will not have purchased the share.” Second, participants answered a series of Likert-type questions assessing a number of dimensions that are potentially relevant to investment decisions (“How much do you like these companies?”; “How much do you trust these companies?” and “How much do you think these companies stand for a good investment?”). Each question was answered on a 5-point scale ranging from “not at all” to “very much”.

Finally, participants indicated their risk aversion by deciding on 10 binary choices (Holt & Laury, 2002; for example: “Which of the following two investment options would you choose from each pair?”, e.g., “240 $ with 10 % probability & 192 $ with 90 % probability” vs. “462 $ with 10 % probability & 21 $ with 90 % probability”. Participants further indicated their behavioural inhibition (7 items, Cronbach’s alpha = .69) and approach style (13 items, Cronbach’s alpha = .79) on a 4-point-Likert scale ranging from “strongly disagree” to “strongly agree” using the BIS-BAS scale (Carver & White, 1994). Data on participants’ risk aversion as well as their behavioural inhibition and approach style was left un-analysed. The complete online survey can be found in the supplemental online material (https://osf.io/7tqj3/).

#### Laboratory

The laboratory experiment consisted of five consecutive parts: (1) preparation for and starting of neurophysiological recording, (2) instructions on the stock broker game, (3) a learning phase in which participants got familiar with the stock characteristics and the stock broker game in general, (4) a paper and pencil test in which participants showed that they had understood the stock characteristics relative to pre-defined criteria, and (5) an investing phase in which participants played the stock broker game.

#### Stock broker game

In the stock broker game, participants chose between pairs of stock options from the eight companies introduced during the online survey. The general task was to make investment decisions by choosing one stock from each pair. Participants were informed that each decision would add one share of that stock to their portfolio, and that they would build eight portfolios throughout the study. As an incentive, participants were told that they would receive a bonus payment based on the value of one randomly chosen portfolio after completing the study^1^. Laboratory instructions for participants can be found in the supplemental online material (https://osf.io/yt4d2/).

The available stock options varied along two objective dimensions: gain and chance. Gain defined the monetary reward obtained if the stock paid out, and was expressed as varying levels of arbitrary units (range: 2.63-11.55). Participants were informed at the start of the study that one unit equalled 2 cents. The chance dimension determined the probability that a given stock would pay out (range: 20%-88%). This manipulation created four different types of stocks, that varied across three levels of expected return (i.e., gain x chance). Stocks were selected so that they would be familiar to Canadian participants, and two companies were used from four market sectors: technology, banking, supermarkets, and motor manufacturing.

The technology companies, Apple and Google, had a high chance of winning a large amount of money (high chance/ high gain; high expected share value: ∼10 units). This meant that the technology companies were the best possible investment for any stock pairing. The two financial sector companies, TD Bank and Scotiabank, had a high chance of winning a small amount of money (high chance/ low gain; medium expected share value: ∼2.3 units). The two retail companies were Sobeys and Loblaws, and had a low chance of winning a large amount of money (low chance/ high gain; medium expected share value: ∼2.3 units). It is important to note that while the supermarkets and banks differed on the levels of gain and chance, they had identical expected monetary value (gain x chance). As such, there is no economic reason why investing in the banks would be a better strategy than investing in the supermarkets. Finally, the motor companies were Ford and Chevrolet, and were characterized as stocks that have a low chance of paying out a small amount of money (low chance/low gain; low expected share value: ∼0.5 units). As such, the motor companies are always the least valuable option in any stock pairing. In total, we tried to align the provided stock characteristics with general subjective representations and real world valuations of the different companies and sectors. The rationale behind this decision was to first of all add high (low) subjective value to high (low) objective conflict. Secondly, we tried to increase the game’s ecologic validity and tangibility by doing so. So while we counterbalanced everything in the stock-broker game we could, each company had the same characteristics for all participants. The exact stock characteristics are depicted in table 1.

**Table 1.**
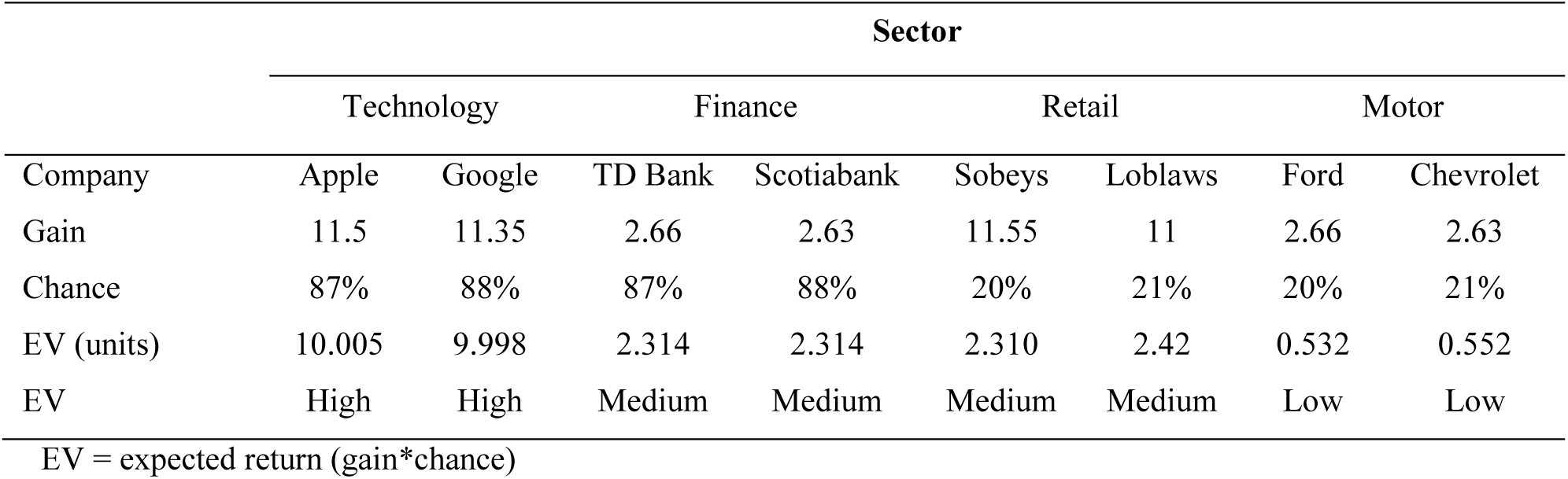
The characteristics of the eight different stocks

#### Learning phase

The study commenced with a learning phase so that participants got familiar with the stock broker game and to ensure that participants learned the stock characteristics to predefined criteria. Two stock options were presented on each trial, and the image of each stock contained the company’s logo and name, in addition to information on gain and chance. When presented on the computer screen, each pair measured approximately 500 x 236 pixels. Stock options from the same sector (i.e., technology, finance, retail, motor manufacturing) were always presented with the same background colour (red, green, purple, and cyan, respectively). These background colours were fully counterbalanced between participants. Targets were presented until response or for a maximum of 2000 ms. Participants were instructed to press the key “f” to choose the option presented on the left side, and to press the key “j” to choose the option presented on the right side of the screen. Very fast responses (RTs < 150 ms) were excluded from further analyses, and responses made faster than the minimum RT and slower than the maximum were followed by “too fast” or “too slow” feedback for 2000 ms, respectively.

The general task was to decide, like a stock-broker, which stock to invest on each trial. A central fixation cross (learning: 37×38 pixels, investing: 19×19 pixels; random duration: 250–750 ms) preceded the target pictures. Consequently, the response-to-stimulus interval varied randomly between 250 and 1,250 ms. The learning phase consisted of 40 trials in which only choices with a clear best option were presented. As such, no conflicting options (i.e., trials where both stocks have equal expected return) were presented to participants during learning. During the learning phase participants also received feedback on the investment outcome following their decision (duration: 1500 ms), indicating whether the stock option they had chosen would have performed well or poorly, and whether they would have earned (or missed) a high or low gain. The feedback was probabilistic and determined based on the selected stock’s chance characteristics. For example, each time an individual invested in Apple, they had an 88% chance of receiving immediate feedback indicating a large gain. Thus, by following the feedback participants were able to learn the characteristics of the stocks over time. Figure 2 provides an overview on decisions in the learning phase.

**Figure 2.**
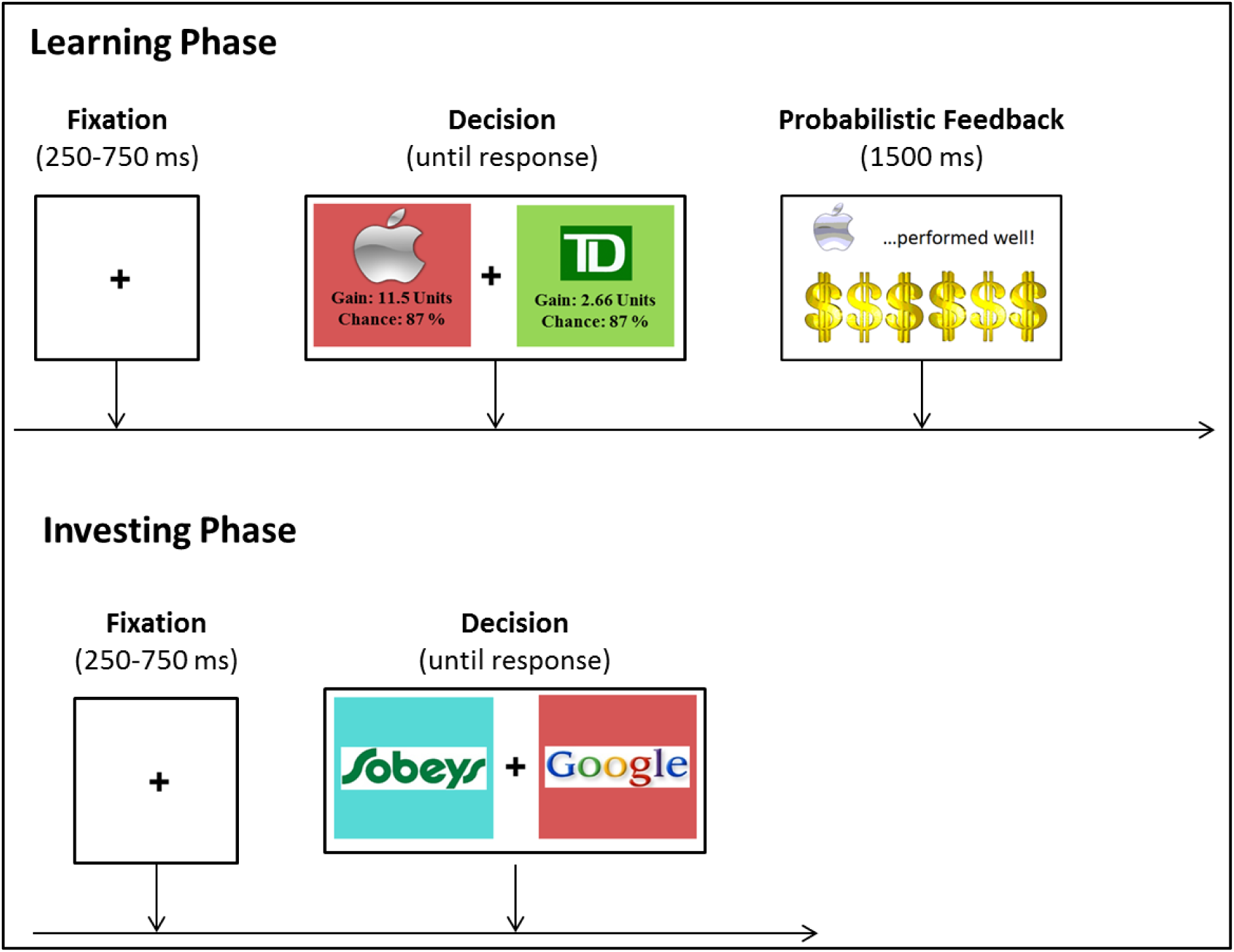
Trial structure in the in the learning (top) and investing (bottom) phase.

#### Paper and pencil test

Before moving beyond the learning phase, participants conducted a paper and pencil test to ensure that they had learned the characteristics of the stock to pre-defined criterion levels. First, participants were asked to indicate which stocks were of low/high gain as well as low/high chance. Second, we asked participants to provide approximate values for the level of gain and chance for each option. Participants did not pass the test if the approximate numbers they gave exceeded +/- 2 units for gain estimates or +/- 10 percentage points chance. If participants did not reach this pre-determined criterion for the initial learning phase, the learning process was repeated again until participants passed the test or completed 4 learning blocks. On average, participants took 2 learning phases to reach criterion.

#### Investing phase

In the final part of the study, participants were instructed that they would now start to make investment decisions that will determine their final profit. Participants were instructed that they would compose eight different portfolios (i.e., groups of 40 sequential investment decisions), out of which one portfolio would be randomly chosen at the end to determine their final profit. For technical reasons, participants always received CAD 5.20 in profit, which was CAD 1 above the maximum expected return of each portfolio.

Throughout the investing phase, we removed the written stock characteristics from the experimental trials and only kept company logos overlaid on the background colours from the original learning phase. This omission ensured that participants made decisions based on both the objective characteristics and their own subjective impressions (based on the online portion of the study). Each pair of images was approximately 192×87 pixels on the computer screen. The experimental period consisted of eight portfolios (i.e., blocks) each containing 40 decisions (i.e., trials), resulting in a total number of 320 trials. In each round, participants were presented with 20 conflicting and 20 non-conflicting pairs of stock options. Conflicting decisions occurred when participants decided between two options with (almost) identical expected monetary value (i.e., Apple vs. Google, TD Bank vs. Scotiabank, Sobeys vs. Loblaws, Ford vs. Chevrolet, TD Bank/Scotiabank vs. Sobeys/Loblaws). Non-conflicting decisions occurred when there was a discrepancy in expected monetary value between stocks (i.e., Apple/Google vs. TD Bank/Scotiabank, Apple/Google vs. Sobeys/Loblaws, Apple/Google vs. Ford/Chevrolet, TD Bank/Scotiabank vs. Ford/Chevrolet, Sobeys/Loblaws vs. Ford/Chevrolet). On a behavioural level we measured participants’ reaction times and percentage choices. With regards to reaction times, choices where reaction times were below 100ms were excluded from all analyses. For percentage choice, we calculated how often participants preferred one stock option over the other. A score of 0 means that participants were fully undecided and that both stocks were chosen equally likely by the participant. A score of 1 means that one stock was always preferred over the other and that the participant was fully decided. Mathematically, percentage choice was defined as: |0.5 - (Frequency of choosing 1 option over the other/ Number of choices)|* 2. Figure 2 provides an overview on decisions in the investing phase.

After every portfolio, participants were asked to rate their feelings when being offered the respective choices by entering a number from 1 (not at all) to 10 (absolutely). We alternated ratings on positive and anxious feelings so that we asked about participants’ positive feelings after four portfolios, and about their anxious feelings after the remaining four portfolios.

### EEG recording

Electroencephalographic (EEG) activity was recorded from 11 Ag/AgCl sintered electrodes embedded in a stretch-lycra cap. The scalp-electrode montage consisted of midline electrode sites (FPz, Fz, FCz, Cz, CPz, Pz & Oz) referenced to the average activity recorded at bilateral earlobes. We made the decision to record primarily from midline electrode sites because the CN is maximal at these sites, and we had no plans of source localizing the electrical signals. Vertical electro-oculography (VEOG) was monitored using a supra-to sub-orbital bipolar montage surrounding the right eye. During recording impedances were monitored (< 5 KΩ) and the EEG signal was digitized at 1024 Hz using ASA acquisition hardware (Advanced Neuro Technology, Enschede, the Netherlands).

Offline the data were band-pass filtered (0.1 to 15 Hz) and corrected for eye-blinks using regression-based procedures (c.f., Gratton, Coles & Donchin, 1983). Automatic procedures were employed to detect and reject EEG artefacts. The criteria applied were a voltage step of more than 25 µV between sample points, a voltage difference of 150 µV within 200 ms intervals, voltages above 85 µV and below -85 µV, and a maximum voltage difference of less than 0.05 µV within 100 ms intervals. These intervals were rejected on an individual channel basis to maximize data retention. All offline EEG data analyses were conducted using Brain Vision Analyzer (v.2.0; Brain Products, GmbH, Gilching, Germany).

For the response locked ERPs, the continuous EEG was segmented into epochs that commenced 500 ms before the response and lasted for 1100 ms. Response-locked ERPs were averaged separately for each choice type and were baseline corrected using a 100 ms window that started 150 ms before the response. We did not use difference scores (e.g., high conflict vs. low conflict) to operationalise the CN. The CN was then operationalized using the mean amplitude in a window 0 to 100 ms after the response at electrodes FCz and Cz. The time-window for the CN was determined using a collapsed localiser method (c.f., Luck & Gaspelin). First, we collapsed over levels of expected value of the best option to generate a grand average ERP across all conditions in the experiment. We then chose a time-window that captured the negative going waveform immediately following the response at electrodes FCz and Cz. ERPs were averaged separately for each pair of stock options (e.g., Apple vs. Google, TD Bank vs. Ford, Sobeys vs. Google, and so on).

### Data analyses

Based on our hypotheses, we divided analyses into three sequential steps. First, we assessed the effects of objective value characteristics (i.e., objective value conflict and amount of expected value) on our dependent measures (hypothesis 1 & 2). In a second step, we tested if response locked ERPs (CN) predict conflict effects on our behavioural and affective DVs (hypothesis 3). In a third step, we tested if subjective impressions (i.e., pre-rated evaluations of each company) change results on our DVs over and above the objective characteristics (hypothesis 4). We only analysed participants’ reactions during the investing phase as in the learning phase participants still got familiar with the stock broker game in general and the characteristics of the different stocks in particular. To account for the nested structure in our data, we used multi-level models (MLM) as they are less restrictive than repeated-measures ANOVA (Field, 2013, p. 818 ff.). MLM, for example, increases statistical power by allowing random effects, and is well able to handle missing data points. As a measure of local effect sizes in mixed-effects regression modelling, Cohen’s *f* was calculated (cf. Selya, Rose, Dierker, Hedeker, & Mermelstein, 2012; “How can I estimate effect size for mixed models?”). All analyses were performed in Stata 13.1 (StataCorp., 2013).

#### Step 1: conflict as a result of objective value characteristics

In a first step, we tested the influence of objective characteristics of the stock pairs on indecision (mean reaction time and percentage choice), affective responses (positive and anxious feelings), and ERPs (CN). We used MLM with the main effects of conflict (high conflict = 1; low conflict = 0) and the expected return of the best option (∼0.5 units = low vs. ∼2.3 units = medium vs. ∼10 units = high), as well as the interaction between these two main effects. High conflict was defined as occurring when each stock in a pair had an (almost) equal expected return (e.g., Google vs. Apple), whereas low conflict pairs had an obvious difference in expected return between stock pairs (e.g., Scotiabank vs. Ford). The interaction term with the expected return of the best option then allows us to test if conflicts scale with the value of the stocks. For example, do high value conflicts (e.g., Google vs. Apple) elicit more conflict reactions than low value conflicts (e.g., Ford vs. Chevrolet). Each MLM had a two-level structure with repeated measures nested within participant. Random intercepts were estimated per participant applying an independent covariance structure. Analyses were performed using the “xtmixed” option in Stata 13.1 (StataCorp., 2013). Simple effect tests were conducted using the “margins” option in Stata 13.1.

#### Step 2: CN amplitude and conflict responses

In a second step, we analysed whether the magnitude of neurophysiological responses to decisions predict the behavioural and affective correlates of conflict. In these analyses the CN was person centered in order to isolate within-subject variance in neural conflict monitoring, and reduce between-subject variability (see Saunders, Milyavskaya, & Inzlicht, 2015, for similar logic). To achieve this we first calculated the mean CN amplitude per participant across all decision types, and then subtracted this from the raw CN score for each of the 28 decision types (e.g., Sobeys vs Ford *minus* participant’s mean CN amplitude). Using this variable—rather than the raw CN amplitude per decision—means that we predict each dependent measure (e.g., positive affect) from within-subject variance in neural conflict monitoring. In other words, this analysis allows us to determine whether within-participant fluctuations in neural conflict monitoring co-vary with fluctuations in within-person positive affect. A key benefit of this analysis strategy is that our sample is well powered to investigate within-subject variation.

To constrain these analyses further, we only tested dependent variables for which we found a conflict effect in the previously defined steps. We ran multilevel analyses where repeated measures were clustered in participants investigating if within-participant fluctuations in CN amplitude predicted behaviour and affect. Analyses were performed using the “xtmixed” option in Stata 13.1 (StataCorp., 2013).

#### Step 3: conflict as a result of subjective value characteristics

Last, we controlled for participants’ subjective impression of the eight companies—measured by the online pre-test—on the previously defined dependent measures. To test subjective influences, we first used confirmatory factor analysis to form a latent variable that reflected participants’ subjective impressions separately for each of the companies. The participants’ ratings (liking, trust, good investment) on the eight companies and the assigned values from the BDM auction to generate a latent factor that estimates how positively individuals viewed each of the eight companies. Analyses were performed using the “gsem” option in Stata 13.1 to control for the multilevel structure of the data (StataCorp., 2013). All eight item sets on the subjective impression showed a good fit to a single-factor model^2^ and values on latent variables were stored for the main analyses. This created a single subjective impression score for each company, with increasing scores indicating increasing subjective value. We then derived variables from these latent factors to create subjective factors analogous to the main effects included in the analysis of the objective characteristics. Mirroring the conflict main effect, we first calculated absolute difference scores on the subjective values for each choice, to create a subjective value main effect. Here, a score of zero would indicate that two options have an equivalent subjective value (i.e., high subjective conflict), with increasing difference scores indicating that one stock out-values the other (i.e., low subjective conflict). The second main effect coded the subjective value of the best rated item from each pair. Finally, we interacted these two variables to test if subjective conflict scales with increasing subjective value.

We again conducted multilevel analyses where repeated measures were clustered in participants. Random intercepts were estimated per participant applying an independent covariance structure. In addition to the objective decision characteristics (choice conflict, expected return of the best option and the interaction term), we entered subjective decision characteristics (subjective conflict, value of the subjectively rated best option and the interaction term) into multilevel analyses. Analyses were performed using the “xtmixed” option in Stata 13.1 (StataCorp., 2013).

## Results

### Conflict reactions as a result of objective value characteristics

We first analysed participants’ behavioural reactions (mean reaction time, percentage choice), reported feelings (positive and anxious feelings) and neurophysiological responses (CN and FCz) in relation to the objective characteristics of the stock task: objective conflict, the expected return of the best option, and the interaction term of these two variables. Table 2 provides an overview of the results.

**Table 2.**
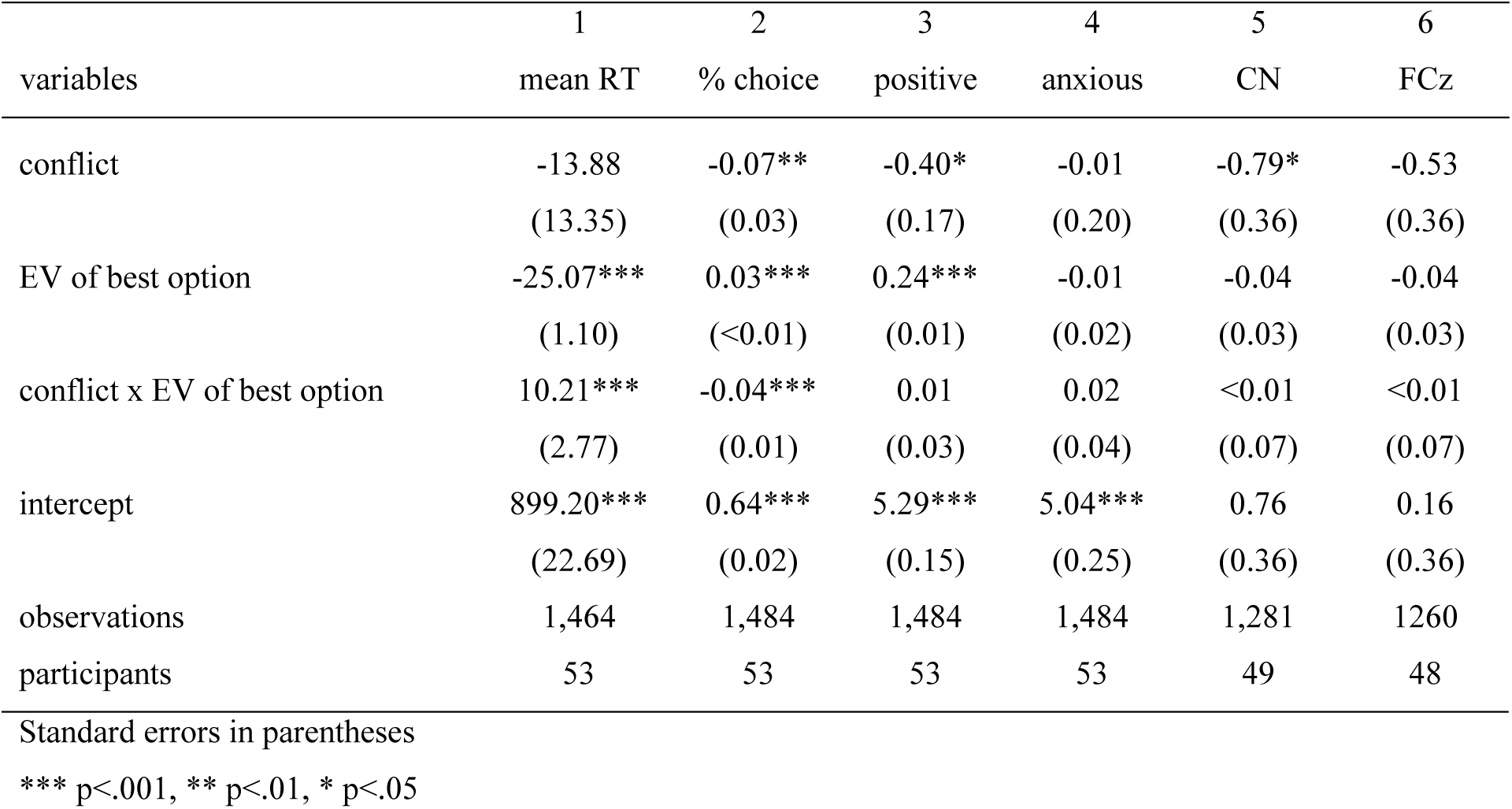
Multilevel regression analyses testing participants’ reactions to objective choice characteristics: objective value conflict, expected value of the best option and their interaction.

#### Mean reaction time

There was no significant main effect of conflict on mean reaction times (*b*=-13.88, *S.E.*=13.35), z=-1.04, *p=*.298, *f^2^* < .001. However, mean reaction times became faster as the expected return of the best option increased (*b*=-25.07, *S.E.*=1.10), *z*=-22.88, *p* < .001, *f^2^* = .371. Though not hypothesized, this can likely be explained as a motivational effect, where people react more quickly to obtain rewards. This effect was less strong for high conflict trials, as indicated by the significant interaction term of conflict with the expected value of the best option, (*b=*10.21, *S.E.=*2.77), z=3.68, *p<*.001, *f^2^* = .009. Altogether, there was a significant effect of conflict on mean reaction time for high EV options, *Chi^2^=*20.00, *p<*.001, but not for low EV options, *Chi^2^=*0.30, *p=*.582, indicating that high value conflict leads to slower decisions.

The results from reaction times support our first and second hypothesis in that participants did become slower in their decisions when facing similarly good options of high expected value.

#### Percentage choice

Results on percentage choices showed a significant main effect of conflict, (*b=*-0.07, *S.E.=*0.03), z=-2.67, *p=*.008, *f^2^* = .004, indicating that participants were more undecided when having to choose between options with equal expected returns (M=54 %, SD = 34 %, 90% CI [0%, 100%]) than options with different expected returns (M=82 %, SD = 28 %, 90% CI [25%, 100%]). As indicated by the significant main effect of the expected value of the best option (*b=*0.03, *S.E.<*0.01), z=11.72, *p<*.001, *f^2^* = .096, participants were generally more decided when more money was at stake. However, a significant interaction effect between conflict and expected value of the best option, (*b=*-0.04, *S.E.=*0.01), z=-7.03, *p*<.001, *f^2^* = .035 and post-hoc tests revealed that the conflict effect was stronger for high EV options, *Chi^2^=*141.69, *p<*.001 than for low EV options, *Chi^2^=*50.88, *p<*.001, suggesting that participants were less decided between options of equal value when more value was at stake.

Taken together, the results from percentage choice are in support of our first and second hypothesis: participants are less decided when facing similarly good options, especially when expected returns of both options are high.

#### Positive and anxious feelings

As would be expected, higher expected returns of the better option led to higher positive feelings (*b=*0.24, *S.E.=*0.01), z=17.24, *p<*.001, *f^2^* = .208. This finding suggests that participants report increasingly positive evaluations of options with increasingly value. Furthermore, participants in general reported slightly lower levels of positive affect for conflicting compared to non-conflicting decisions, (*b=*-0.40, *S.E.=*0.17), z=-2.37, *p=*.018, *f^2^* = .004, yet, this effect did not become stronger with increasing values of the expected value of the best option, (interaction term: *b=*0.01, *S.E.=*0.03), z=0.38, *p=*.706, *f^2^* < .001.

Regarding anxious feelings, higher expected returns of the better option did not lead to less anxious feelings (*b=*-0.01, *S.E.=*0.02), z=-0.36, *p=*.720, *f^2^* < .001. Furthermore, we did not find that participants reported higher levels of anxious feelings for conflicting compared to non-conflicting decisions, (*b=*-0.01, *S.E.=*0.20), z=-0.03, *p=*.974, *f^2^* < .001. Nor did we find an interaction effect between conflict and the expected value of the best option, (*b=*0.02, *S.E.=*0.04), z=0.62, *p=*.535, *f^2^* < .001.

Taken together, the results on reported feelings suggest that high conflict trials lead to slightly lower positive feelings, while the expected return of the better option increases positive affect. However, in contrast to past results (e.g., Shenhav & Buckner, 2014), we do not observe any effects on felt anxiety levels. Based on these results we derive that conflict is aversive (hypothesis 1) in the sense that at least the absence of conflict is pleasing.

Figure 3 (top panels) illustrates the results on reported feelings as a function of objective conflict.

**Figure 3.**
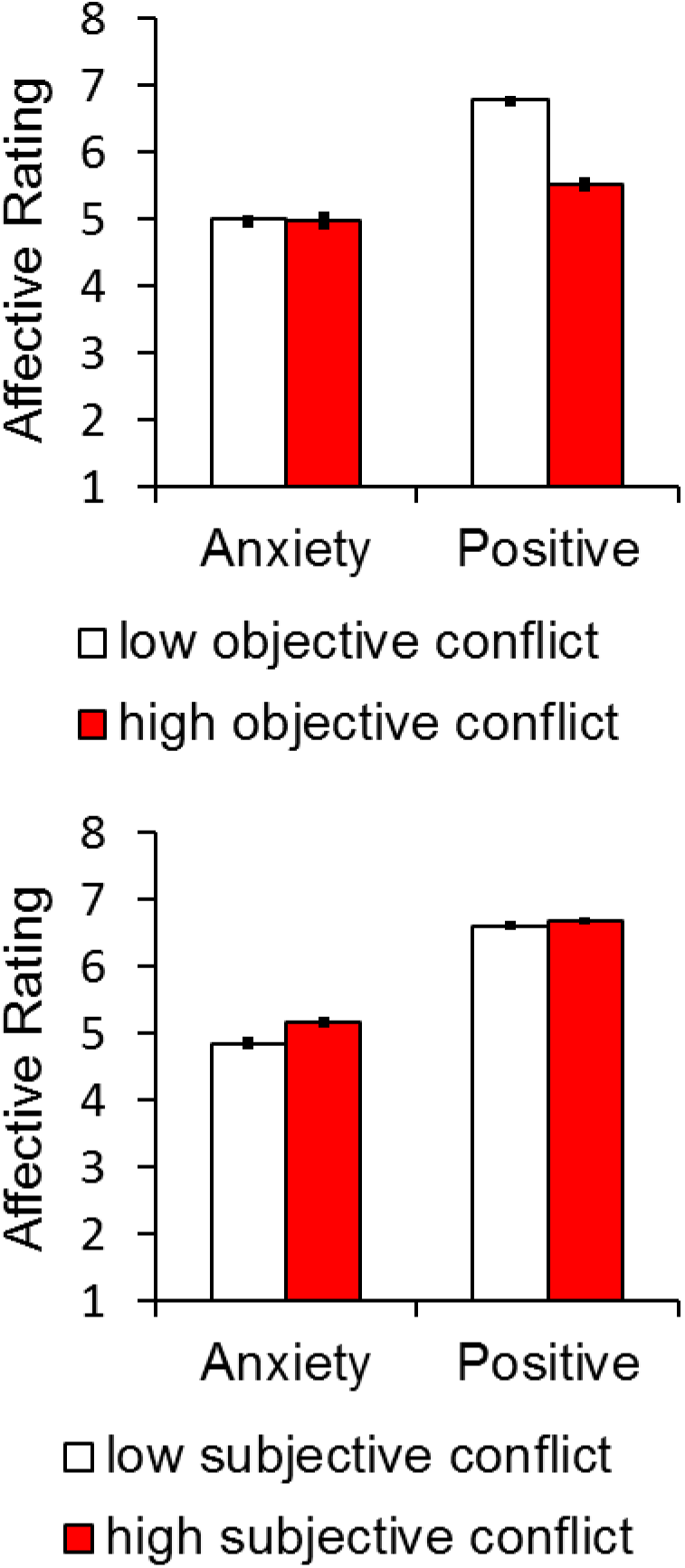
Bar charts showing the effects of objective conflict on reported levels of anxiety and positive affect (top panels) and the independent effect of subjective conflict (difference in subjective impression) on anxiety and conflict. Low and high subjective conflict reflect plus or minus 1 SD from mean on the difference in subjective value between stock options, respectively (lower panels). Estimated means are presented in the bottom panel for display purposes only. Error bars reflect within-subject 95% confidence intervals.

#### CN

A significant main effect of conflict on CN was found (*b=*-0.79, *S.E.=*0.36), z=- 2.21, *p=*.027, *f^2^* = .004^3^, at electrode site Cz indicating that high conflict trials induce slightly more conflict-related negativity than low conflict trials, see figure 4. Expected returns of the best option did not have any influence on CN (*b=*-0.4, *S.E.=*0.03), z=-1.35, *p=*0.178, *f^2^* = .002, nor did the interaction of expected return of the best option with conflict (*b<*0.01, *S.E.=*0.07), z=0.05, *p=*0.959, *f^2^* < .001.

**Figure 4.**
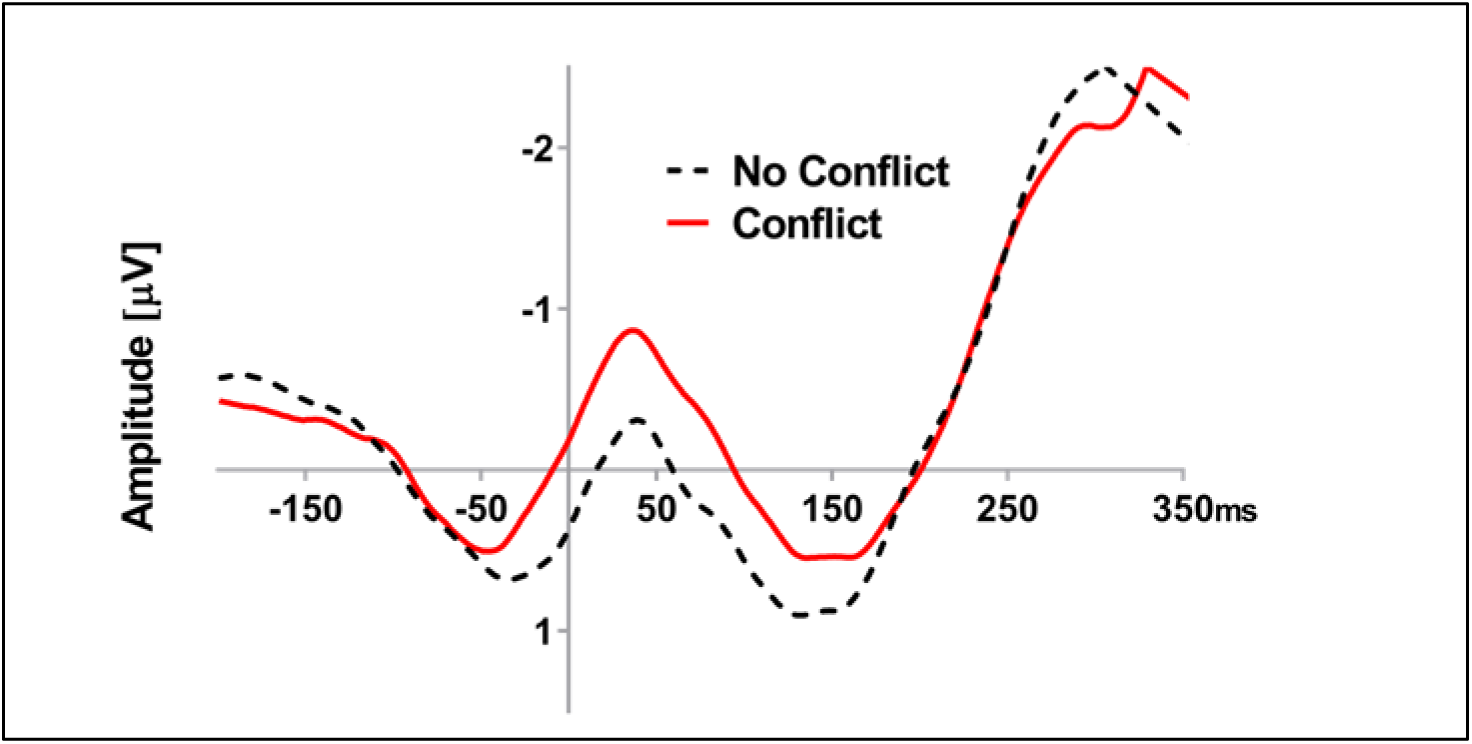
ERP waveforms showing effect on conflict on the CN at electrode Cz.

Though modest, the results are in-line with our hypothesis 1 and reveal that deciding between two equally good options induces higher neurophysiological conflict reactions at electrode site Cz in participants. However, contrary to hypothesis 2 and the observed behavioural effects on reaction times and percentage choice, this effect did not scale with increasing expected returns.

#### FCz

With regards to electrode site FCz we did not find any main or interaction effects: conflict had no main effect on FCz (*b=*-0.53, *S.E.=*0.36), z=-1.47, *p=*.142, *f^2^* = .001, neither did expected returns of the best option (*b=*-0.04, *S.E.=*0.03), z=-1.35, *p=*0.178, *f^2^* = .001, nor interaction of expected return of the best option with conflict (*b<*0.01, *S.E.=*0.07), z=-0.05, *p=*0.961, *f^2^* < .001.

### CN amplitude and conflict reactions

With regards to our third hypothesis, we were interested in how the magnitude of neural reactivity to conflict will go hand-in-hand with behavioural and emotional reactions to conflict. We therefore tested how the behavioural and affective correlates of conflict and thus positive feelings and percentage choice (see table 2), are associated with CN amplitude. All trials (high and low conflict) are considered in this set of analyses. As analyses on electrode FCz did not reveal an effect of conflict, we did not test for behavioural and affective correlates with electrode FCz.

#### CN and positive feelings

The CN predicted positive feelings on a 10% alpha level (*b=*0.03, *S.E.=*0.02), z=1.86, *p=*.062, *f^2^* = .047, with modest effect sizes, providing a first indication that higher neurophysiological conflict reactivity was somewhat accompanied by reduced positive affect to a decision type. The CN results on reported feelings hence indicate that neurophysiological conflict reactivity during financial decisions is modestly associated with participants’ positive feelings.

#### CN and percentage choice

Running multilevel analyses with the CN predicting participants’ choices, we found that the CN significantly predicted participants’ choices (*b=*0.01, *S.E.<*0.01), z=4.33, *p<*.001, *f^2^* = .026, indicating that higher neurophysiological conflict reactivity during choices was associated with increased indecision. Though the effect size is modest, the results indicate that neurophysiological responses directly after financial decisions correlate to participants’ indecision.

### Conflict reactions as a result of subjective value characteristics

In a third step, we tested if participants also pull value information from subjective sources. In all analyses, the effects of objective conflict, value of the objectively best option and their interaction stayed qualitatively the same when including our subjective measures, unless stated otherwise. Table 3 provides an overview of all results on this set of analyses.

**Table 3.** Multilevel regression analyses testing participants’ reactions to subjective value characteristics in addition to objective choice characteristics

| variables | 1<br>RT | 2<br>%choice | 3<br>positive | 4<br>anxious | 5<br>CN | 6<br>FCz |
| --- | --- | --- | --- | --- | --- | --- |
| conflict | -8.40<br>(13.72) | -0.07*<br>(0.03) | -0.44*<br>(0.18) | -0.02<br>(0.21) | -0.70<br>(0.38) | -0.45<br>(0.38) |
| EV of best option | -24.93***<br>(1.15) | 0.03***<br>(<0.01) | 0.25***<br>(0.02) | -0.01<br>(0.02) | -0.04<br>(0.03) | -0.05<br>((0.03) |
| conflict x EV of best option | 9.64***<br>(2.79) | -0.04***<br>(0.01) | 0.02<br>(0.04) | 0.02<br>(0.04) | -0.01<br>(0.08) | -0.01<br>(0.08) |
| subjective conflict | -5.51<br>(10.16) | 0.02<br>(0.02) | 0.19<br>(0.13) | 0.35*<br>(0.16) | 0.27<br>(0.29) | 0.17<br>(0.29) |
| value of subj. best option | -27.01*<br>(12.13) | 0.02<br>(0.02) | 0.40**<br>(0.16) | -0.12<br>(0.19) | 0.11<br>(0.34) | 0.03<br>(0.34) |
| subj. conflict x subj. best option | 6.86<br>(9.70) | -0.02<br>(0.02) | -0.13<br>(0.13) | -0.10<br>(0.15) | -0.36<br>(0.27) | -0.27<br>(0.34) |
| intercept | 902.90***<br>(25.14) | 0.61***<br>(0.03) | 5.03***<br>(0.18) | 4.92***<br>(0.28) | 0.65<br>(0.43) | 0.17<br>(0.43) |
| observations | 1,344 | 1,344 | 1,344 | 1,344 | 1,141 | 1120 |
| participants | 48 | 48 | 48 | 48 | 44 | 43 |

Standard errors in parentheses
\*\*\* $p < .001$ , \*\* $p < .01$ , \* $p < .05$

#### Mean reaction time

Adding subjective conflict measures to multilevel analyses, we found that a higher value of the subjectively rated best option made participants decide even faster (*b=*-27.01, *S.E.=*12.13), z=-2.23, *p=*.026, *f^2^* = .004. Though the effect size is very small, the results suggest that value of the subjectively rated best option goes into the same direction as the expected value of the objectively best option. No effects on reaction times were found for neither subjective conflict (*b=*-5.50, *S.E.=*10.16), z=-0.54, *p=*.558, *f^2^* < .001, nor the interaction of subjective conflict and subjectively rated best option (*b=*6.86, *S.E.=*9.69), z=0.71, *p=*.480, *f^2^* < .001. Effects resulting from the subjectively rated best option seem to point into the same direction as the effects from the objective counterpart that is the expected value of the best option.

#### Percentage choice

No effects on percentage choice were found for subjective conflict (*b=*0.02, *S.E.=*0.02), z=1.20, *p=*.232, *f^2^* < .001, the subjectively rated best option (*b=*0.02, *S.E.=*0.02), z=0.80, *p=*.427, *f^2^* < .001, or the interaction of subjective conflict and subjectively rated best option (*b=*-0.02, *S.E.=*0.02), z=-1.00, *p=*.319, *f^2^* < .001.

#### Positive and anxious feelings

Subjective value affected positive feelings: a higher value of the subjectively rated best option went along with slightly higher positive feelings (*b*=0.40, *S.E.*=0.16), z=2.59, *p=*.010, *f^2^* = .004. However, we found no effects on positive feelings for subjective conflict (*b=*0.19, *S.E.=*0.13), z=1.44, *p=*.150, *f^2^* = .002, or the interaction of subjective conflict with subjectively rated best option (*b=*-0.13, *S.E.=*0.13), z=- 1.04, *p=*.299, *f^2^* < .001.

With regards to feelings of anxiety, we found that conflict based on subjective ratings slightly increased anxious feelings in participants (*b*=0.35, *S.E.*=0.16), z=2.23, *p=*.025, *f^2^* = .004. No effects were found for the subjectively rated best option (*b=*-0.12, *S.E.=*0.19), z=- 0.65, *p=*.514, *f^2^* < .001, or the interaction of subjective conflict and the subjectively rated best option (*b=*-0.10, *S.E.=*0.15), z=-0.64, *p=*.522, *f^2^* < .001, on felt anxiety. Subjective conflict was the only independent variable that significantly influenced felt anxiety and felt anxiety was the only dependent variable reacting to subjective value conflict. Figure 3 (lower panels) illustrates the results on reported feelings as a function of subjective conflict.

#### CN

We found no effects on CN for either subjective conflict (*b=*0.27, *S.E.=*0.29), z=0.94, *p=*.348, *f^2^* < .001, the subjectively rated best option (*b=*0.11, *S.E.=*0.34), z=0.33, *p=*.741, *f^2^* < .001, or the interaction of subjective conflict and subjectively rated best option (*b=*-0.36, *S.E.=*0.27), z=-1.37, *p=*.172, *f^2^* = .002. Including subjective conflict measures to multilevel analyses, the objective conflict effect on CN only remained significant at a 10% alpha level (*b=*-0.70, *S.E.=*0.38), z=-1.82, *p=*.069, *f^2^* = .003. **FCz.** We found no effects on FCz for either subjective conflict (*b=*0.17, *S.E.=*0.29), z=0.58, *p=*.562, *f^2^* < .001, the subjectively rated best option (*b=*0.03, *S.E.=*0.34), z=0.08 *p=*.939, *f^2^* < .001, or the interaction of subjective conflict and subjectively rated best option (*b=*-0.27, *S.E.=*0.27), z=-1.00, *p=*.318, *f^2^* = .001.

## Discussion

We investigated behavioural reactions, reported feelings, and neurophysiological responses to incentivized financial decision conflict. Our results are at odds with classic economic models. From a purely rational perspective (cf. Schoemaker, 1982; Fehr & Hoff, 2011), facing objectively equal options should not challenge individuals, meaning that choices between equal options should be straightforward. However, this is not what our data suggests. Participants were more undecided and less pleased when they decided between options of equal value compared to decision scenarios in which one option clearly outperformed the other. Moreover, the amplitude of the CN, a negative-going ERP that is thought to represent decision conflict in the medial prefrontal cortex (Di Domenico et al., 2016), was more negative on conflicting decisions relative to non-conflicting decisions. Regarding the monetary value of decision options, behavioural reactions became even stronger when more money was at stake: Participants were even more undecided and also slower in decisions when they deliberated between conflicting options of high monetary value. Contrary to classic economic theories, our results suggest that it is particularly challenging to decide between equivalent rewards, and, furthermore, we identify the CN as a neurophysiological signal that tracks this decision conflict.

Our findings support previous psychological and neuroscientific research indicating that decisions between equally valued options trigger conflict at the behavioural and neural level (e.g., Lin et al., 2017; Nakao et al., 2010; Nakao et al., 2013; Shenhav & Buckner, 2014). The results also extend previous research in two important ways. First of all, by applying varying monetary values we can show on a behavioural level (i.e. reaction times and indecision) that conflict reactions scale with value, whereas it appears that the CN amplitude captures reactions to conflict but does not scale with value. Secondly, in previous studies participants decided between qualitatively different things (dancer vs. chemist; ipod vs. sudoku book, small reward now vs. larger reward later). These previous results could have been explained from an economic perspective with the concept of opportunity costs, stating that “the true cost of something is what you give up to get it” (http://www.economist.com/economics-a-to-z/o). In our study decisions were made between options with identical monetary outcomes and conflict was triggered during decisions though participants could not lose anything by design. This means that no opportunity costs were involved by design ruling out this alternative economic explanation.

While our results are at odds with assumptions made by classic economic theory, they might shed light on seemingly anomalous observations from finance research. It is well documented, for example, that private investors show low levels of trading activity (e.g., “Shareownership 2000”, 2002), a phenomenon commonly known as investor inertia. For example, in the domain of retirement savings in the US, private investors largely avoid financial decisions altogether and, as a result, miss opportunities to optimize their portfolios (e.g., Madrian & Shea, 2001; Thaler & Benartzi, 2004). Such inertia might be related to the difficulty that arises during financial decision making across behavioural, affective, and neural levels of analysis.

Our data further reveals that the CN amplitude not only reacts to conflict decisions but also predicts participants’ affective reactions and behavioural responses. Importantly, the association between reductions in positive affect and CN amplitude should be interpreted with some caution (and merits future replication) given that this effect was not significant at our *apriori* alpha level (*p* < .05). Nevertheless and under the assumption that positive and negative affect exist on a continuum (Russel, 2003), these neural results are consistent with recent suggestions that the aMCC not only detects decision conflicts (Botvinick et al., 2001), but also with suggestions that aMCC tracks negatively valenced events during cognitive control (e.g., Botvinick, 2007; Inzlicht, Bartholow & Hirsh, 2015; Koban & Pourtois, 2014; Shackman et al., 2011) and decision making (e.g., Saunders, Lin, Milyavskaya, & Inzlicht, 2017; Shenhav & Buckner, 2013). To our knowledge, the current results are the first to link the CN to indecision and subjective evaluations that arise during decision making.

Consistent with the components name (i.e., the conflict negativity), the CN has typically been associated with conflict. However, it should be considered if the component perhaps reflects processes other than the conflict alone. Foremost, it might be suggested that rather than manipulating conflict, our task manipulated decision difficulty. In line with recent accounts of the medial prefrontal cortex (Shenhav, Straccia, Cohen & Botvinick, 2014) it is possible that the CN is tracking decision difficulty rather than conflict. However, it can be particularly difficult to dissociate effort from conflict. In many psychological and cognitive neuroscience models, conflict is proposed to trigger a range of remedial cognitive adjustments including changes in attention and adjustments in performance (Botvinick et al., 2001; Proulx et al., 2012). These considerations have led authors to suggest that conflict typically coincides with an aversive and effortful state (Botvinick, 2007). As such, high-conflict trials in our paradigm likely co-occurred with an effortful state in which individuals deliberated between equally valued options.

### Subjective preference and investment decisions

Recent findings from real-world trading data suggest that investors do not only consider objective stock characteristics, but additionally discriminate based on subjective impressions resulting from, for example, social information about ethnicity or gender (Kumar, et al., 2015; Niessen-Ruenzi & Ruenzi, 2015). Based on these findings we explored how subjective preferences might impact conflict decisions. Contrary to our expectations, subjective influences on decision conflict were not particularly strong. Still, our study mildly supports the suggestion that idiosyncratic preferences impact investment decisions over and above objective stock characteristics (i.e., gain and risk). Participants reacted faster and reported more positive feelings about their decisions the higher they subjectively valued one company over the other; and this was beyond the objective characteristics of the stocks. Subjective impressions were based on participants’ actual brand perception of the companies assessed on average six days before the participants entered the experiment. Our results hence support the idea that private investors are mildly influenced by subjective sources of information in addition to objective sources. This is particularly surprising in the current context, where participants were trained explicitly and to criterion on objective stock characteristics.

### Limitations and future research

Despite conceptually similar designs, our affect results do not fully mirror those of Shenhav and Buckner (2014). In this earlier study, subjective conflict between desirable consumer goods (e.g., digital camera vs. camcorder) produced simultaneous increases in anxiety *and* positive affect relative to less conflicting decisions (e.g., iPod vs. a Sudoku book). In our study, however, participants felt more anxious but not more positive towards subjective value conflicts, whereas objective conflict only reduced positive affect. While many differences exist between these studies (e.g., consumer goods vs. financial decision making; fMRI vs. EEG; trial numbers and timing, the learning element in the current investigation), we suggest that the difference in results might be best accounted for by differences in overall value involved for each decision (e.g., Kachelmeier & Shehata, 1992), with absolute values being significantly lower in our study.

The effect of conflict on the CN was also less statistically robust than is typically observed for other similar frontocentral ERPs, and was only found at electrode Cz but not FCz. The ERN and FRN, for example, show sizeable and remarkably robust modulations in response to both the accuracy of responses and the valence of externally provided feedback (Holroyd & Coles, 2002; Yeung et al., 2005). As such, our results should be considered a preliminary investigation into the effects of conflict during financial decision making on this ERP. One potential source unaccounted for error in the current study, one that perhaps contributes to the size of the observed effects, is the idiosyncratic variability in value judgements during subjective choice. Perhaps conflict in the domain of subjective value can be more robustly modulated if the choice scenarios are tailored to each individual’s subjective preferences. Indeed, we have recently demonstrated robust modulation of the CN to conflict when decision stimuli were created on an individual-by-individual bases using earlier subjective preferences indicated by the participants (Lin et al., 2017). Thus, while the results are consistent across studies, it is possible that idiographic paradigms elicit particularly robust effects on the conflict negativity.

As already stated above, we find on a behavioural level that reactions to conflict scale with value. At the same time we observe in our data that the CN amplitude and positive feelings capture conflict reactions but these conflict reactions do not scale with value. This interesting pattern should be further addressed by future research, especially, if the CN really mainly detects the mere existence of conflict.

Our results are generative for future ERP research exploring decision making across multiple domains with increasing ecological validity. While some of the evidence reported in the current study should be interpreted with some caution given the exploratory nature of our work, in addition to the high p-values for some effects (e.g., the association between positive affect and CN amplitude), the current methodology provides a paradigm that can be used to test conflicts that more closely resemble the decision we make in our day to day lives across multiple domains in ERP experiments. It is our hope that readers of this work will adopt such methods not only to explore the neural correlates of conflict in the laboratory, but also how these constrained neural reactions predict real-world outcomes.

### Conclusion

We examined how conflict derived from objective and subjective value characteristics of stocks affect investment decisions and their neural correlates. We demonstrated that in a financial context, decisions and affective reactions are influenced by subjective and objective value conflict. Moreover, we provide novel evidence that the CN serves as a neural correlate to objective value conflict and correlates with behavioural indecision. Our key finding is that choosing in a situation where it should not matter which option to pick from an economic perspective, alerts the CN to the extent that investors are more undecided. Our results, thus, may serve as one out of several indicators as to why private investors avoid financial decisions and remain with suboptimal portfolio allocations over time.

## Author Note

Gesa-K. Petersen and Blair Saunders contributed equally to this article. This research was supported in part by a grant from the Excellence Initiative of the German Research Foundation (DFG).

The authors do not have any interests that might be interpreted as influencing the research.

Correspondence concerning this article should be addressed to Gesa-Kristina Petersen, Munich School of Management, Schackstrasse 4, 80539 Munich, Germany.

## Footnotes

1 Due to technical reasons, all participants were paid CAD 5.20 at the end of the study, which was CAD 1 above the expected return of each portfolio if the participant had always chosen the economically best available option.

2 Apple (4 items, *χ ^2^*=0.85, *p=*.655; CFI=1.00, TLI=1.06, RMSEA<.01 [90% CI=.00, .215]), Google (4 items, *χ ^2^*=0.11, *p*=.095; CFI=1.00, TLI=1.26, RMSEA<.01 [90% CI=.00, .04]), TD Bank (4 items, *χ ^2^*=0.78, *p*=.675; CFI=1.00, TLI=1.09, RMSEA<.01 [90% CI=.00, .21]), Scotiabank (4 items, *χ ^2^*=10.20, *p*=.006; CFI=.88, TLI=0.65, RMSEA=.28 [90% CI=.13, .47]), Loblaws (4 items, *χ ^2^*=2.00, *p*=.367; CFI=1.00 TLI=1.00, RMSEA<.01 [90% CI=.00, .28]), Sobeys (4 items, *χ ^2^*=1.57, *p*=.456; CFI=1.00 TLI=1.04, RMSEA<.01 [90% CI=.00, .26]), Chevrolet (4 items, *χ ^2^*=13.35, *p*=.001; CFI=0.77, TLI=0.32, RMSEA=.33 [90% CI=.18, .51]) and Ford (4 items, *χ ^2^*=1.61, *p*=.448; CFI=1.00 TLI=1.02, RMSEA<.01 [90% CI=.00, .26]).

3 As a first indicator of motor response effects we additionally tested, if effects get affected by reaction times. However, conflict effects on CN do not qualitatively change when we additionally control for reaction times

## References

1. Anderson, C. J. (2003). The psychology of doing nothing: Forms of decision avoidance result from reason and emotion. Psychological Bulletin, 129(1), 139–167. https://doi.org/10.1037/0033-2909.129.1.139

2. Aristotle, (1922). On the Heavens (Trans. J.L. Stocks). Retrieved from http://www.sacred-texts.com/cla/ari/oth/index.htm (Original work published 350 BC), book II, part 13, 295b

3. Bartholow, B. D., Pearson, M. A., Dickter, C. L., Sher, K. J., Fabiani, M., & Gratton, G. (2005). Strategic control and medial frontal negativity: Beyond errors and response conflict. Psychophysiology, 42(1), 33–42. https://doi.org/10.1111/j.1469-8986.2005.00258.x

4. Becker, G. M., DeGroot, M. H., & Marschak, J. (1964). Measuring utility by a single-response sequential method. Behavioral Science, 9(3), 226–232. https://doi.org/10.1002/bs.3830090304

5. Botvinick, M. M. (2007). Conflict monitoring and decision making: Reconciling two perspectives on anterior cingulate function. Cognitive, Affective, & Behavioral Neuroscience, 7, 356–366. https://doi.org/10.3758/cabn.7.4.356

6. Botvinick, M. M., Braver, T. S., Barch, D. M., Carter, C. S., & Cohen, J. D. (2001). Conflict monitoring and cognitive control. Psychological Review, 108(3), 624–652. https://doi.org/10.1037/0033-295x.108.3.624

7. Brown, J. W., & Braver, T. S. (2005). Learned predictions of error likelihood in the anterior cingulate cortex. Science, 307, 1118–1121.

8. Carver, C. S., & White, T. L. (1994). Behavioral inhibition, behavioral activation, and affective responses to impending reward and punishment: The BIS/BAS Scales. Journal of Personality and Social Psychology, 67(2), 319–333. https://doi.org/10.1037/0022-3514.67.2.319

9. Cavanagh, J. F., Figueroa, C. M., Cohen, M. X., & Frank, M. J. (2012a). Frontal theta reflects uncertainty and unexpectedness during exploration and exploitation. Cerebral Cortex, 22, 2575–2586.

10. Cavanagh, J. F., & Frank, M. J. (2014). Frontal theta as a mechanism for cognitive control. Trends in Cognitive Sciences, 18, 414–421.

11. Cavanagh, J. F., Zambrano-Vazquez, L., & Allen, J. J. (2012b). Theta lingua franca: A common mid-frontal substrate for action monitoring processes. Psychophysiology, 49, 220–238. https://doi.org/10.1111/j.1469-8986.2011.01293.x

12. Di Domenico, S. I., Le, A., Liu, Y., Ayaz, H., & Fournier, M. A. (2016). Basic psychological needs and neurophysiological responsiveness to decisional conflict: an event-related potential study of integrative self-processes. Cognitive, Affective, & Behavioral Neuroscience, 16(5), 848–865. https://doi.org/10.3758/s13415-016-0436-1

13. Faul, F., Erdfelder, E., Lang, A. G., & Buchner, A. (2007). G* Power 3: A flexible statistical power analysis program for the social, behavioral, and biomedical sciences. Behavior Research Methods, 39(2), 175–191. https://doi.org/10.3758/bf03193146

14. Fehr, E., & Hoff, K. (2011). Introduction: Tastes, castes and culture: The influence of society on preferences. The Economic Journal, 121(556), F396–F412. https://doi.org/10.1111/j.1468-0297.2011.02478.x

15. Festinger, L. (1962). A theory of cognitive dissonance (Vol. 2). Stanford University Press.

16. Field, A. (2013). Discovering statistics using IBM SPSS statistics. Sage.

17. Gratton, G., Coles, M. G., & Donchin, E. (1983). A new method for off-line removal of ocular artifact. Electroencephalography and Clinical Neurophysiology, 55(4), 468–484. https://doi.org/10.1016/0013-4694(83)90135-9

18. Hare, T. A., Camerer, C. F., & Rangel, A. (2009). Self-control in decision-making involves modulation of the vmPFC valuation system. Science, 324(5927), 646–648. https://doi.org/10.1126/science.1168450

19. Holroyd, C. B., & Coles, M. G. (2002). The neural basis of human error processing: Reinforcement learning, dopamine, and the error-related negativity. Psychological Review, 109(4), 679. http://dx.doi.org/10.1037/0033-295X.109.4.679

20. Holt, C. A., & Laury, S. K. (2002). Risk aversion and incentive effects. American Economic Review, 92(5), 1644–1655. https://doi.org/10.1257/000282802762024700

21. How can I estimate effect size for mixed models? Stata FAQ. UCLA: Institute for digital research and education. Retrieved from https://stats.idre.ucla.edu/stata/faq/how-can-i-estimate-effect-size-for-mixed/.

22. Inzlicht, M., Bartholow, B. D., & Hirsh, J. B. (2015). Emotional foundations of cognitive control. Trends in Cognitive Sciences, 19(3), 126–132. https://doi.org/10.1016/j.tics.2015.01.004

23. Johnson, E. J., & Goldstein, D. (2003). Do defaults save lives? Science, 302(5649), 1338–1339. https://doi.org/10.1126/science.1091721

24. Kachelmeier, S. J., Shehata, M. (1992). Examining risk preferences under high monetary incentives: Experimental evidence from the people’s republic of china. American Economic Review, 82(5), 1120–1141. http://www.jstor.org/stable/2117470

25. Kerns, J. G., Cohen, J. D., MacDonald, A. W., Cho, R. Y., Stenger, V. A., & Carter, C. S. (2004). Anterior cingulate conflict monitoring and adjustments in control. Science, 303, 1023–1026.

26. Koban, L., & Pourtois, G. (2014). Brain systems underlying the affective and social monitoring of actions: an integrative review. Neuroscience and Biobehavioral Reviews, 46(1), 71–84. https://doi.org/10.1016/j.neubiorev.2014.02.014

27. Kumar, A., Niessen-Ruenzi, A., & Spalt, O. G. (2015). What’s in a name? Mutual fund flows when managers have foreign-sounding names. Review of Financial Studies, 28(8), 2281–2321. https://doi.org/10.1093/rfs/hhv017

28. Lin, H., Saunders, B., Hutcherson, C. A., & Inzlicht, M. (2017). Midfrontal theta and pupil dilation parametrically track subjective conflict (but also surprise) during intertemporal choice. NeuroImage.

29. Madrian, B. C., & Shea, D. F. (2001). The power of suggestion: Inertia in 401 (k) participation and savings behavior. The Quarterly Journal of Economics, 116(4), 1149–1187. https://doi.org/10.1162/003355301753265543

30. Nakao, T., Bai, Y., Nashiwa, H., & Northoff, G. (2013). Resting-state EEG power predicts conflict-related brain activity in internally guided but not in externally guided decision-making. NeuroImage, 66(1), 9–21. https://doi.org/10.1016/j.neuroimage.2012.10.034

31. Nakao, T., Mitsumoto, M., Nashiwa, H., Takamura, M., Tokunaga, S., Miyatani, M., … & Watanabe, Y. (2010). Self-knowledge reduces conflict by biasing one of plural possible answers. Personality and Social Psychology Bulletin, 36(4), 455–469. https://doi.org/10.1177/0146167210363403

32. Niessen-Ruenzi, A., & Ruenzi, S. (2015). Sex matters: Gender bias in the mutual fund industry. https://doi.org/10.2139/ssrn.1957317

33. Proulx, T., Inzlicht, M., & Harmon-Jones, E. (2012). Understanding all inconsistency compensation as a palliative response to violated expectations. Trends in Cognitive Sciences, 16(5), 285–291. https://doi.org/10.1016/j.tics.2012.04.002

34. Richard, R., Van der Pligt, J., & De Vries, N. (1996). Anticipated affect and behavioral choice. Basic and Applied Social Psychology, 18(2), 111–129. https://doi.org/10.1002/(sici)1099-0771(199609)9:3<185::aid-bdm228>3.0.co;2-5

35. Saunders, B., Lin, H., Milyavskaya, M., & Inzlicht, M. (2017). The emotive nature of conflict monitoring in the medial prefrontal cortex. International Journal of Psychophysiology. https://doi.org/10.1016/j.ijpsycho.2017.01.004

36. Saunders, B., Milyavskaya, M., & Inzlicht, M. (2015). What does cognitive control feel like? Effective and ineffective cognitive control is associated with divergent phenomenology. Psychophysiology, 52(9), 1205–1217. https://doi.org/10.1111/psyp.12454

37. Schoemaker, P. J. (1982). The expected utility model: Its variants, purposes, evidence and limitations. Journal of Economic Literature, 20(2), 529–563. http://www.jstor.org/stable/2724488

38. Schwarz, N. (2000). Emotion, cognition, and decision making. Cognition and Emotion, 14(4), 433–440. https://doi.org/10.1080/026999300402745

39. Selya, A. S., Rose, J. S., Dierker, L. C., Hedeker, D., & Mermelstein, R. J. (2012). A practical guide to calculating Cohen’s f2, a measure of local effect size, from PROC MIXED. Frontiers in Psychology, 3, 111. https://doi.org/10.3389/fpsyg.2012.00111

40. Shackman, A. J., Salomons, T. V., Slagter, H. A., Fox, A. S., Winter, J. J., & Davidson, R. J. (2011). The integration of negative affect, pain and cognitive control in the cingulate cortex. Nature Reviews Neuroscience, 12(3), 154–167. https://doi.org/10.1038/nrn2994

41. Shareownership 2000 Highlights. (2002). Retrieved from http://www.nyxdata.com/nysedata/asp/factbook/viewer_edition.asp?mode=text&key=51&category=11

42. Shenhav, A., & Buckner, R. L. (2014). Neural correlates of dueling affective reactions to win–win choices. Proceedings of the National Academy of Sciences, 111(30), 10978–10983. https://doi.org/10.1037/abn0000196

43. Shenhav, A., Straccia, M. A., Cohen, J. D., & Botvinick, M. M. (2014). Anterior cingulate engagement in a foraging context reflects choice difficulty, not foraging value. Nature Neuroscience, 17, 1249–1254.

44. StataCorp. 2013. Stata Statistical Software: Release 13. College Station, TX: StataCorp LP.

45. Thaler, R. H., & Benartzi, S. (2004). Save more tomorrow™: Using behavioral economics to increase employee saving. Journal of Political Economy, 112(S1), 164–187. https://doi.org/10.1086/380085

46. Vidal, F., Hasbroucq, T., Grapperon, J., & Bonnet, M. (2000). Is the ‘error negativity’ specific to errors? Biological Psychology, 51(2), 109–128. https://doi.org/10.1016/S0301-0511(99)00032-0

47. Yeung, N., Botvinick, M. M., & Cohen, J. D. (2004). The neural basis of error detection: conflict monitoring and the error-related negativity. Psychological Review, 111(4), 931. http://dx.doi.org/10.1037/0033-295X.111.4.931

48. Zeelenberg, M. (1999). Anticipated regret, expected feedback and behavioral decision making. Journal of Behavioral Decision Making, 12(2), 93–106. https://doi.org/10.1002/(sici)1099-0771(199906)12:2<93::aid-bdm311>3.0.co;2-s

